# Multiplex genome engineering in *Clostridium beijerinckii* NCIMB 8052 using CRISPR-Cas12a

**DOI:** 10.1101/2022.08.22.504755

**Authors:** Constantinos Patinios, Stijn T. de Vries, Mamou Diallo, Lucrezia Lanza, Pepijn L. J. V. Q. Verbrugge, Ana M. López-Contreras, John van der Oost, Ruud A. Weusthuis, Servé W. M. Kengen

**Author notes:** These authors have contributed equally to the work (Constantinos Patinios, Stijn T. de Vries, Mamou Diallo). Corresponding author: Servé W. M. Kengen.

## Abstract

*Clostridium* species are re-emerging as biotechnological workhorses for industrial acetone-butanol-ethanol production. This re-emergence is largely due to advances in fermentation technologies but also due to advances in genome engineering and re-programming of the native metabolism. Several genome engineering techniques have been developed including the development of several CRISPR-Cas tools. Here, we expanded the CRISPR-Cas toolbox and developed a CRISPR-Cas12a genome engineering tool in *Clostridium beijerinckii* NCIMB 8052. By controlling the expression of FnCas12a with the strict xylose-inducible promoter, we achieved efficient (25-100%) single-gene knockout of five *C. beijerinckii* NCIMB 8052 genes (*Spo0A, Upp, Cbei_1291, Cbei_3238, Cbei_3832*). Moreover, we achieved multiplex genome engineering by simultaneously knocking out the *Spo0A* and *Upp* genes in a single step with an efficiency of 18%. Finally, we showed that the spacer sequence and position in the CRISPR array can affect the editing efficiency outcome.

## Introduction

*Clostridium beijerinckii* NCIMB 8052, a gram-positive, spore-forming, anaerobic bacterium, is a member of the acetone-butanol-ethanol (ABE) producing *Clostridium* species. ABE fermentation has a considerable industrial history as it played a major role during the twentieth century and especially during World War I for the production of acetone (to produce cordite/gunpowder), butanol, lacquer solvents and jet fuel, before being outcompeted by petrochemical processes (1-4). Due to economic and environmental reasons and due to advances in biotechnology, sustainable ABE fermentation using *Clostridium* species has re-emerged (5).

To take advantage of the industrial potential of *Clostridium* species, several genome engineering tools have been developed (6). The genome engineering tools can be broadly divided into the ones which rely on homologous recombination (HR) based allelic exchange and the ones which depend on group II intron-retargeting mutagenesis (TargeTron/ClosTron) (7,8). Whilst the group II intron-retargeting mutagenesis generally allows quick and efficient genome engineering, it has several disadvantages including the inability to target genes smaller than 400 bp, it interrupts rather than deletes the target of interest, it frequently relies on the genomic integration of antibiotic resistance genes and the intron may be spliced back out by its associated intron-encoding protein (9). On the other hand, HR-based techniques enable the generation of scarless and complete deletion mutants and they do not rely on the integration of antibiotics in the genome of the target organism. For high accuracy, however, HR should be combined with an efficient counterselection mechanism such as CRISPR-Cas (10). Following this requirement, CRISPR-Cas in combination with HR has been used to achieve high editing efficiencies in various *Clostridia* species as summarized by McAllister and Song (2019).

To date, several CRISPR-Cas-based tools have been developed for *C. beijerinckii* (9,11-16). Most of the developed CRISPR-Cas tools rely on the Cas9 nuclease and only a small fraction is based on the Cas12a nuclease. The preference towards Cas9 is probably due to a first-comer effect and due to its successful application in various organisms (17). However, Cas12a has distinctive advantageous features over Cas9 including smaller size (Cas12a: ∼1300 a.a., Cas9: ∼1600 a.a.) and recognition of a T-rich 5’-(T)TTV-3’ PAM site at the 5’ end of the protospacer sequence which increases the number of target sites in AT-rich organisms like *Clostridia* (∼30% GC-content) (18). In addition, Cas12a can process its own crRNA array due to its RNase activity; a feature that makes Cas12a ideal for multiplex genome engineering as the transcription of a single CRISPR array expressed by a single promoter is the only requirement for the generation of multiple crRNAs (18).

In this study, we used the FnCas12a nuclease to create single- and multi-gene deletions in *C. beijerinckii* NCIMB 8052. Depending on the target gene, single-gene knockout efficiencies varied from 25% to 100%. Multiplex (two gene) deletion was also achieved in one step with a knockout efficiency of 18%. Spacer sequence and position in the CRISPR array affected the multiplex knockout efficiency, revealing potential limitations and room for improvement.

## Materials and methods

### Microbial strains and growth conditions

Table 1 shows all the strains used or generated in this study. *Escherichia coli* NEB^®^ 5-alpha was used for plasmid assembly and cloning following the manufacturer’s instructions (NEB). Transformed *E. coli* cells were grown at 37°C in LB liquid medium (10 g L^-1^ tryptone, 5 g L^-1^ yeast extract, 10 g L^-1^ NaCl) or on LB agar plates (LB liquid medium, 15 g L^-1^ bacteriological agar) containing spectinomycin (0.1 g L^-1^).

**Table 1.** Strains used or generated in this study.

| Strain | Genotype | Source |
| --- | --- | --- |
| *Escherichia coli* NEB® 5-alpha | *fhuA2 Δ(argF-lacZ)U169 phoA glnV44 Φ80 Δ(lacZ)M15 gyrA96 recA1 relA1 endA1 thi-1 hsdR17* | NEB |
| *Clostridium beijerinckii* NCIMB 8052 | Wild type | (20) |
| *C. beijerinckii* NCIMB 8052<br>$\Delta Spo0A$ | $\Delta Spo0A$ (*Cbei_1712*) | This study |
| *C. beijerinckii* NCIMB 8052<br>$\Delta Upp$ | $\Delta Upp$ (*Cbei_0408*) | This study |
| *C. beijerinckii* NCIMB 8052<br>$\Delta Spo0A$, $\Delta Upp$ | $\Delta Spo0A$ (*Cbei_1712*), $\Delta Upp$ (*Cbei_0408*) | This study |
| *C. beijerinckii* NCIMB 8052,<br>$\Delta Cbei_1291$ | $\Delta Cbei_1291$ | This study |
| *C. beijerinckii* NCIMB 8052,<br>$\Delta Cbei_3238$ | $\Delta Cbei_3238$ | This study |
| *C. beijerinckii* NCIMB 8052,<br>$\Delta Cbei_3932$ | $\Delta Cbei_3932$ | This study |

Transformed *C. beijerinckii* NCIMB 8052 cells were grown anaerobically at 37°C in modified clostridial growth medium containing glucose as the main carbon source (mCGM-G: 5 g L^-1^ yeast extract, 0.75 g L^-1^ KH_2_PO_4_, 0.75 g L^-1^ K_2_HPO_4_, 0.4 g L^-1^ MgSO_4_. 7H_2_O, 0.01 g L^-1^ MnSO_4_. H_2_O, 0.01 g L^-1^ FeSO_4_. 7H_2_O, 1 g L^-1^ NaCl, 2 g L^-1^ L-asparagine, 2 g L^-1^ (NH_4_)_2_SO_4_, 0.125 g L^-1^ L-cysteine, 13.753 g L^-1^ D-(+)-glucose. H_2_O) or on mCGM-G agar (1 g L^-1^ yeast extract, 2 g L^-1^ tryptone, 0.5 g L^-1^ KH_2_PO_4_, 1 g L^-1^ K_2_HPO_4_, 0.1 g L^-1^ MgSO_4_. 7 H_2_O, 0.01 g L^-1^ MnSO_4_. H_2_O, 0.015 g L^-1^ FeSO_4_. 7 H_2_O, 0.01 g L^-1^ CaCl_2_, 0.002 g L^-1^ CoCl_2_, 0.002 g L^-1^ ZnSO_4_, 2 g L^-1^ (NH_4_)_2_SO_4_, 55 g L^-1^ D-(+)-glucose. H_2_O, 12 g L^-1^ agar) supplemented with 0.65 g L^-1^ spectinomycin.

For fermentation assays and knockout generation, *C. beijerinckii* NCIMB 8052 cells were grown in GAPES medium (2.5 g L^-1^ yeast extract, 1 g L^-1^ KH_2_PO_4_, 0.61 g L^-1^ K_2_HPO_4_, 1 g L^-1^ MgSO_4_. 7H_2_O, 0.0066 g L^-1^ FeSO_4_. 7 H_2_O, 2.9 g L^-1^ C_2_H_7_NO_2_, 0.19 g L^-1^ pABA, 0.125 g L^-1^ L-cysteine, 65.6 g L^-1^ D-(+)-glucose. H_2_O) supplemented with 0.65 g L^-1^ spectinomycin (19).

To induce the expression of *FnCas12a*, transformed *C. beijerinckii* NCIMB 8052 cells were plated on mCGM-X agar (containing 40 g L^-1^ xylose instead of glucose as the carbon source) supplemented with 0.65 g L^-1^ spectinomycin.

### Plasmid construction and transformation

The plasmids used in this study are shown in Table 2. Unless otherwise specified, all plasmids were assembled through NEBuilder^®^ HiFi DNA Assembly (NEB). The basic backbone plasmid pCOMA_NT-crRNA was constructed by amplifying the pCB102 ori, colE1 ori and aad9 from pWUR100S (pS), FnCas12a from pY002 and the XylR-XylBP from pE_X_cas9. The crRNA was ordered as synthetic gene fragment (Twist Bioscience).

**Table 2.** Plasmids used in this study.

| Plasmid name | Relevant characteristics | Reference |
| --- | --- | --- |
| pY002 | p15A ori, Lacp-WT FnCas12a, TetR, CmR | (18) |
| pWUR100S (pS) | pCB102 ori, colE1 ori, Aad9R | (23) |
| pE_X_cas9 | colE1 ori, pAMβ1 ori, 2 µ ori, AmpR, ErmR, URA3 | (23) |
| pCOMA_NT-crRNA | pCB102 ori, colE1 ori, ThlP-NT-crRNA-ThlT, XylRP-XylR, XylBP-FnCas12a-FdxT, Aad9R | This study |
| pCOMA_Spo0A-crRNA | pCB102 ori, colE1 ori, ThlP-Spo0A-crRNA-ThlT, XylRP-XylR, XylBP-FnCas12a-FdxT, Aad9R | This study |
| pCOMA_NT-crRNA_Spo0AHA | pCB102 ori, colE1 ori, ThlP-NT-crRNA-ThlT, XylRP-XylR, XylBP-FnCas12a-FdxT, Aad9R, 500 bp homologous arms for Spo0A | This study |
| pCOMA_Spo0A-crRNA_Spo0AHA | pCB102 ori, colE1 ori, ThlP-Spo0A-crRNA-ThlT, XylRP-XylR, XylBP-FnCas12a-FdxT, Aad9R, 500 bp homologous arms for Spo0A | This study |
| pCOMA_NT-crRNA_UppHA | pCB102 ori, colE1 ori, ThlP-NT-crRNA-ThlT, XylRP-XylR, XylBP-FnCas12a-FdxT, Aad9R, 500 bp homologous arms for Upp | This study |
| pCOMA_Upp1-crRNA_UppHA | pCB102 ori, colE1 ori, ThlP-Upp1-crRNA-ThlT, XylRP-XylR, XylBP-FnCas12a-FdxT, Aad9R, 500 bp homologous arms for Upp | This study |
| pCOMA_Cbei_1291-crRNA_Cbei_1291HA | pCB102 ori, colE1 ori, ThlP- Cbei_1291-crRNA-ThlT, XylRP-XylR, XylBP-FnCas12a-FdxT, Aad9R, 500 bp homologous arms for Cbei_1291 | This study |
| pCOMA_ Cbei_3238-crRNA_Cbei_3238HA | pCB102 ori, colE1 ori, ThlP- Cbei_3238-crRNA-ThlT, XylRP-XylR, XylBP-FnCas12a-FdxT, Aad9R, 500 bp homologous arms for Cbei_3238 | This study |
| pCOMA_ Cbei_3932-crRNA_Cbei_3932HA | pCB102 ori, colE1 ori, ThlP- Cbei_3932-crRNA-ThlT, XylRP-XylR, XylBP-FnCas12a-FdxT, Aad9R, 500 bp homologous arms for Cbei_3932 | This study |
| pCOMA_NT-crRNA_Spo0AHA_UppHA | pCB102 ori, colE1 ori, ThlP-NT-crRNA-ThlT, XylRP-XylR, XylBP-FnCas12a-FdxT, Aad9R, 500 bp homologous arms for Spo0A, 500 bp homologous arms for Upp | This study |
| pCOMA_Spo0A-Upp1-crRNA_Spo0AHA_UppHA | pCB102 ori, colE1 ori, ThlP-Spo0A-Upp1-crRNA-ThlT, XylRP-XylR, XylBP-FnCas12a-FdxT, Aad9R, 500 bp homologous arms for Spo0A, 500 bp homologous arms for Upp | This study |
| pCOMA_Upp1-Spo0A-crRNA_Spo0AHA_UppHA | pCB102 ori, colE1 ori, ThlP-Upp1-Spo0A-crRNA-ThlT, XylRP-XylR, XylBP-FnCas12a-FdxT, Aad9R, 500 bp homologous arms for Spo0A, 500 bp homologous arms for Upp | This study |
| pCOMA_Spo0A-Upp2-crRNA_Spo0AHA_UppHA | pCB102 ori, colE1 ori, ThlP-Spo0A-Upp2-crRNA-ThlT, XylRP-XylR, XylBP-FnCas12a-FdxT, Aad9R, 500 bp homologous arms for Spo0A, 500 bp homologous arms for Upp | This study |
| pCOMA_Upp2-Spo0A-crRNA_Spo0AHA_UppH A | pCB102 ori, colE1 ori, ThlP-Upp2-Spo0A-crRNA-ThlT, XylRP-XylR, XylBP-FnCas12a-FdxT, Aad9R, 500 bp homologous arms for Spo0A, 500 bp homologous arms for Upp | This study |

To introduce the homology arms into the pCOMA_NT-crRNA plasmid series, 1 μg pCOMA_NT-crRNA was linearized using AccI (NEB). The linear backbone was then dephosphorylated using shrimp alkaline phosphatase (rSAP, NEB) following the manufacturer’s instructions. rSAP and residual AccI were deactivated by incubating the solution at 80°C for 20 min. The linearized backbone was further purified using the DNA clean and concentrator kit (Zymo Research). The homology arms were amplified by PCR using *C. beijerinckii* NCIMB 8052 genomic DNA as template and the oligonucleotides listed in Table S1. The correct size of the homology arms was confirmed through gel electrophoresis. Following, the amplicons were gel purified using the zymoclean gel DNA recovery kit (Zymo Research) and introduced into the linearized pCOMA_NT-crRNA through NEBuilder^®^ HiFi DNA Assembly (NEB) following the instructions from the manufacturer. 5 μL of the assembly was used to transform *E. coli* NEB^®^ 5-alpha cells (NEB). Transformed cells were plated on LB agar containing spectinomycin (0.1 g L^-1^) and incubated at 37°C overnight. Obtained colonies were screened through PCR and the obtained plasmids were sequenced for the correctness of the homology arm insert using Sanger sequencing (Macrogen Europe B.V.).

Single or double targeting spacers were introduced through Golden Gate assembly using an adapted protocol (21). Briefly, 1 μL of each of the two complementary oligonucleotides (100 μM each; Table S1) were mixed with 1 μL NaCl (1 M) and 47 μL MQ water and incubated at 95°C for 5 min. Following, the solution was cooled down at room temperature for at least 2 h to achieve annealing of the complementary oligonucleotides. The annealed oligonucleotides were then diluted 10 times and 2 μL of the diluted oligonucleotides was mixed with 2 μL (0.01 - 0.02 pmol μL^-1^) of the appropriate pCOMA_NT-crRNA plasmid series and 2 μl of MetaMix stock (10 μl BsaI-HF®v2, 15 μL T4 ligation buffer, 10 μL T4 ligase and 15 μL MQ). The mix solution was then incubated in a thermocycler using the following protocol: 5 min at 37°C, 5 min at 16°C followed by 5 min at 37°C (repeat for 15-30 cycles), 5 min at 37°C, 20 min at 80°C. 1 μL of the solution was then used to transform chemically competent *E. coli* NEB^®^ 5-alpha (NEB) according to the instructions of the manufacturer. Transformed cells were plated on LB agar plates containing spectinomycin (0.1 g L^-1^) and incubated overnight at 37°C. Obtained colonies were used for plasmid cloning by growing them in 10 mL LB medium containing spectinomycin (0.1 g L^-1^), incubating overnight at 37°C. Plasmid purification was performed by using the GeneJET Plasmid Miniprep kit (ThermoFischer Scientific) and following the manufacturer’s instructions. To verify the spacer(s) sequence, plasmids were sequenced using Sanger sequencing (Macrogen Europe B.V.).

*C. beijerinckii* was transformed as previously described (22). In detail, 100 μL of heat-shocked *C. beijerinckii* spores (1 min at 99°C) were used to inoculate 25 mL of mCGM liquid medium followed by overnight incubation at 37°C. 20 mL of the overnight culture was used to inoculate 180 mL of pre-warmed (37°C) mCGM liquid medium which was then incubated at 37°C until an OD_600_ equal to 0.3-0.4 was reached. Following, in an anaerobic tent, the culture was transferred into a 400-mL sterile centrifuge tube which was then sealed with parafilm to limit oxygen entrance. The culture was then centrifuged aerobically at 6000 rpm at 4°C for 10 min. The centrifuged culture was put on ice and transferred again in the anaerobic tent. The supernatant was then discarded and the cell pellet was resuspended with 25 mL of ice-cold anaerobic electroporation buffer (270 mM D-sucrose, 1 mM sodium phosphate buffer pH 7.4, 1 mM MgCl_2_). The resuspended culture was transferred into a 30-mL sterile centrifuge tube which was then sealed with parafilm to limit oxygen entrance. The resuspended culture was then centrifuged aerobically at 6000 rpm at 4°C for 10 min followed by discarding the supernatant and resuspending the cell pellet with 1.5 mL of ice-cold anaerobic electroporation buffer. 300 μL of the resuspended cells were used to electroporate (1.25 kV, 25 μF, 100 D) 3-5 μg plasmid DNA using 0.2 cm ice-cold electroporation cuvettes. After electroporation, the transformants were recovered at 37°C in 3 mL of anaerobic mCGM for 3-4 h. After recovery, cultures were centrifuged at 6000 rpm for 5 min and the cell pellet was plated on mCGM solid medium containing spectinomycin (0.65 g L^-1^) followed by incubation at 37°C for 72-96 h. Obtained colonies were screened for the presence of FnCas12a, the crRNA and the homology arms through colony PCR. Correct transformants were subcultured in mCGM and stored as vegetative cells in 20% glycerol at -80°C until use.

### *C. beijerinckii* NCIMB 8052 knockout generation

*C. beijerinckii* cells transformed with the pCOMA plasmid series were grown in 25 mL selective GAPES medium for 48-96 h to allow for homologous recombination to occur. 100 μL of the fully grown culture was then plated on selective mCGM-X agar plates to induce expression of *FnCas12a* and allow for the counterselection of the mutants. Agar plates were incubated for 24-48 h at 37°C and obtained colonies were screened through colony PCR. To perform colony PCR, colonies were resuspended in 50 μL PBS buffer (pH 7.4) and heated for 10 min at 99°C. 1 μL of the heated solution was then used as template for PCR using the Q5^®^ High-Fidelity DNA Polymerase (NEB), following the manufacturer’s instructions. The primers used for colony PCR are listed in Table S1. Lastly, to confirm the complete knockout of the gene of interest, PCR amplicons were sequenced through Sanger sequencing (Macrogen). Each knockout experiment was performed in triplicate and the average knockout efficiency was calculated by defining the percentage of clean mutants (i.e., no mixed bands) versus wild type and mix bands.

### Plasmid curing

To cure the *C. beijerinckii* NCIMB 8052 knockout strains of the pCOMA plasmids, the cells were grown in 25 mL mCGM-G liquid medium without antibiotics for 24 h. 100 μL of the grown culture was then plated on mCGM-G agar plates without the presence of antibiotics and grown for 24 h at 37°C. Obtained colonies were randomly selected and streaked out on one mCGM-G agar plate with antibiotics and on one mCGM-G agar plate without antibiotics and incubated for 24 h at 37°C. Colonies that did not grow on selective medium but grew on non-selective medium were selected and screened for the absence of plasmid through colony PCR. Mutant colonies which lost the respective pCOMA plasmid were grown in GAPES medium without antibiotics and glycerol stocks were made and stored at -80°C until further use.

### ΔS*po0a* and WT *C. beijerinckii* NCIMB 8052 fermentation assays and morphology

The WT and Δ*Spo0A C. beijerinckii* NCIMB 8052 strains were grown in GAPES medium without antibiotics for 48 h. At different time intervals, 1 mL of headspace was recovered and the solvent concentration was determined using gas chromatography (GC). 1 mL of liquid culture was also recovered, of which the pH, OD_600_ and organic acid concentration were determined using high-pressure liquid chromatography (HPLC).

A Shimadzu GC-2010 equipped with an Agilent technologies DB-WAX UI GC column (30 m x 0.53 mm) using a temperature gradient of 60-125°C over 10 min and a nitrogen flow rate of 115 mL min^-1^ was used to separate metabolites. A split ratio of 20 and a carrier flow program with a constant pressure of 30 kPa was applied. References of GAPES medium containing 100, 50, 20 and 5 mM of acetone, ethanol and butanol were used to create a calibration curve. As internal standard, 5 mM of 1-propanol was used.

For HPLC, a Shimadzu LC-2030 with a Shimadzu RID-20A detector was used. To separate the metabolites, a Shodex SUGAR SH1821 column was operated at 45°C with a flow rate of 0.8 mL min^-1^ and a flow time of 20 mins. 0.01N H2SO4 was used as eluent. References of 100; 50; 25; 12.5; 6.25; 3.125, 1.5625 and 0.78125 mM of lactate, acetate and butyrate were used to create a calibration curve. As internal standard, 5 mM of crotonate was used.

Pictures of WT and Δ*Spo0A C. beijerinckii* NCIMB 8052 colonies were taken on mCGM agar plates using Carl Zeiss Axio Scope.A1, 100 x total magnification, phase 1.

## Results and discussion

### Markerless deletion of *Spo0a* through inducible expression of FnCas12a

Several CRISPR-Cas tools have been developed for genome engineering of *C. beijerinckii* strains (6,9,11-13,15,16). However, multiplex gene editing in *C. beijerinckii* has never been demonstrated. We sought to develop an easy-to-use and efficient genome editing tool for multiplex genome engineering in the strain *C. beijerinckii* NCIMB 8052 using CRISPR-Cas12a. Cas12a was chosen as the appropriate Cas protein due to its ability to process its CRISPR array and therefore facilitate multiplex gene targeting using the expression of a single crRNA transcript (18).

A previous report showed successful single-gene genome engineering of *C. beijerinckii* NCIMB 8052 using AsCas12a (15). The AsCas12a used in this study was derived from pDEST-hisMBP-AsCpf1-EC (Addgene plasmid #79007) which has a codon optimized nucleotide sequence for *E. coli*. Since *C. beijerinckii* NCIMB 8052 and *E. coli* differ in genomic GC-content (30% versus 51%, respectively), the use of the *E. coli* optimized AsCpf1 in *C. beijerinckii* NCIMB 8052 may have retarded translation speed and fidelity (Fig. S1) (24,25). Considering codon usage as an important factor for successful protein expression and folding, we reasoned that we should use a *Cas12a* gene that matches the codon usage of *C. beijerinckii* NCIMB 8052 and also follows the same translation speed and fidelity (24). To this end, we have chosen the wild type FnCas12a nuclease based on its low GC percentage (30%) and matching codon usage for *C. beijerinckii* NCIMB 8052 (Fig. S1).

To develop a simple genome engineering tool for *C. beijerinckii* NCIMB 8052, a single plasmid approach was used containing the *FnCas12a* gene, the CRISPR array (repeat-spacer-repeat) with an insertion site for easy exchange of the spacer through Golden Gate and a multiple cloning site (MCS) to insert the homology arms and facilitate gene knockout (Fig. 1). To control the expression of FnCas12a and avoid potential cell toxicity (12,26-29) we chose to use the xylose-inducible promoter derived from *C. difficile* (16,30,31). The use of the xylose-inducible system has a dual function in our setup as it can serve both as the inducer molecule for the expression of FnCas12a but also as carbon- and energy-source for growth. Therefore the use of glucose in the growth medium can be omitted and the effect of potential catabolite repression can be avoided (31). To express the crRNA, we chose to use the strong constitutive thiolase promoter (ThlP).

**Figure 1.**
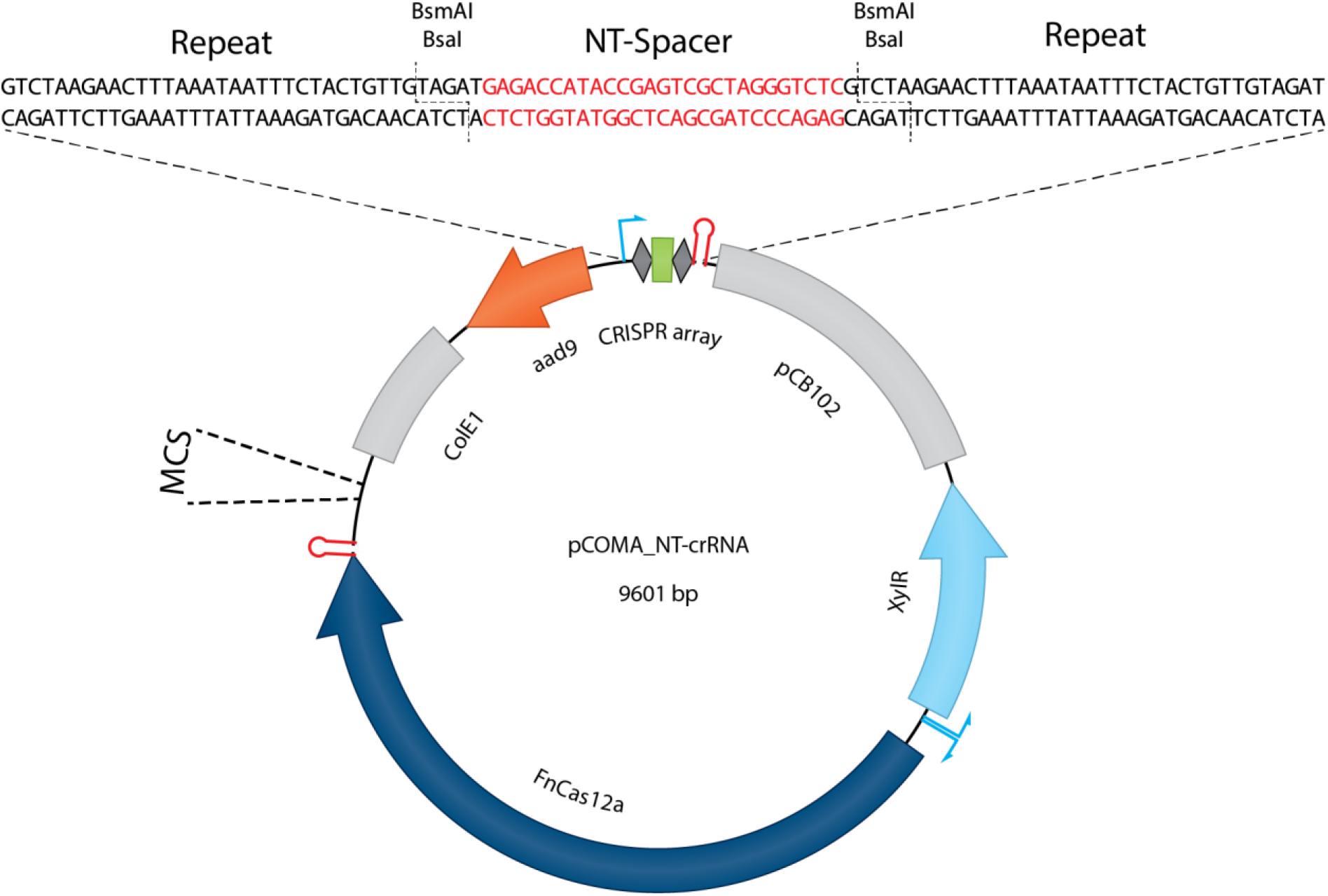
Backbone of the pCOMA plasmid series. At the top of the plasmid the CRISPR array is shown, where a non-targeting (NT) spacer can be conveniently replaced using Golden Gate. Homologous arms can be inserted at the multiple cloning site (MCS) using Gibson Assembly.

As a proof-of-principle, we selected the well-characterized *Spo0A* (Cbei_1712) gene as a knockout target. The Δ*Spo0A* strain has a distinctive morphological and metabolite production phenotype, making it easy to identify *Spo0A* knockouts (32,33). To this end, plasmids pCOMA_NT-crRNA (non-targeting control), pCOMA_Spo0A-crRNA (targeting), pCOMA_NT-crRNA_Spo0AHA (non-targeting control with homology arms) and pCOMA_Spo0A-crRNA_Spo0AHA (targeting with homology arms) were constructed and transformed to *C. beijerinckii* NCIMB 8052 cells. Correct transformants were grown for 48 h in mCGM-G liquid medium, after which 100 μL of the culture were plated on mCGM-G solid medium (as a control) and 100 μL of the culture on mCGM-X solid medium to induce the expression of FnCas12a and, hence, counterselect for mutants.

Transformants plated on mCGM-G solid medium showed comparable numbers of colonies (approximately 10^3^), regardless of the transformed plasmid (Fig. 2A). Similarly, transformants carrying a non-targeting spacer, led to approximately 10^3^ colonies when plated on mCGM-X. In contrast, transformants carrying a targeting spacer and plated on mCGM-X showed fewer colonies compared to the non-targeting controls. As expected, transformation with pCOMA_Spo0A-crRNA resulted in very few colonies (∼10), which shows the functionality of FnCas12a to successfully target and cleave the genome of *C. beijerinckii* NCIMB 8052. The presence of a small number of colonies may be attributed to PAM, spacer, protospacer, crRNA or FnCas12a mutants that could escape the counterselective properties of FnCas12a. A reduced number of colonies (∼51) compared to the non-targeting controls, but 10x higher than pCOMA_Spo0A-crRNA was observed when the pCOMA_Spo0A-crRNA_Spo0AHA plasmid was used.

**Figure 2.**
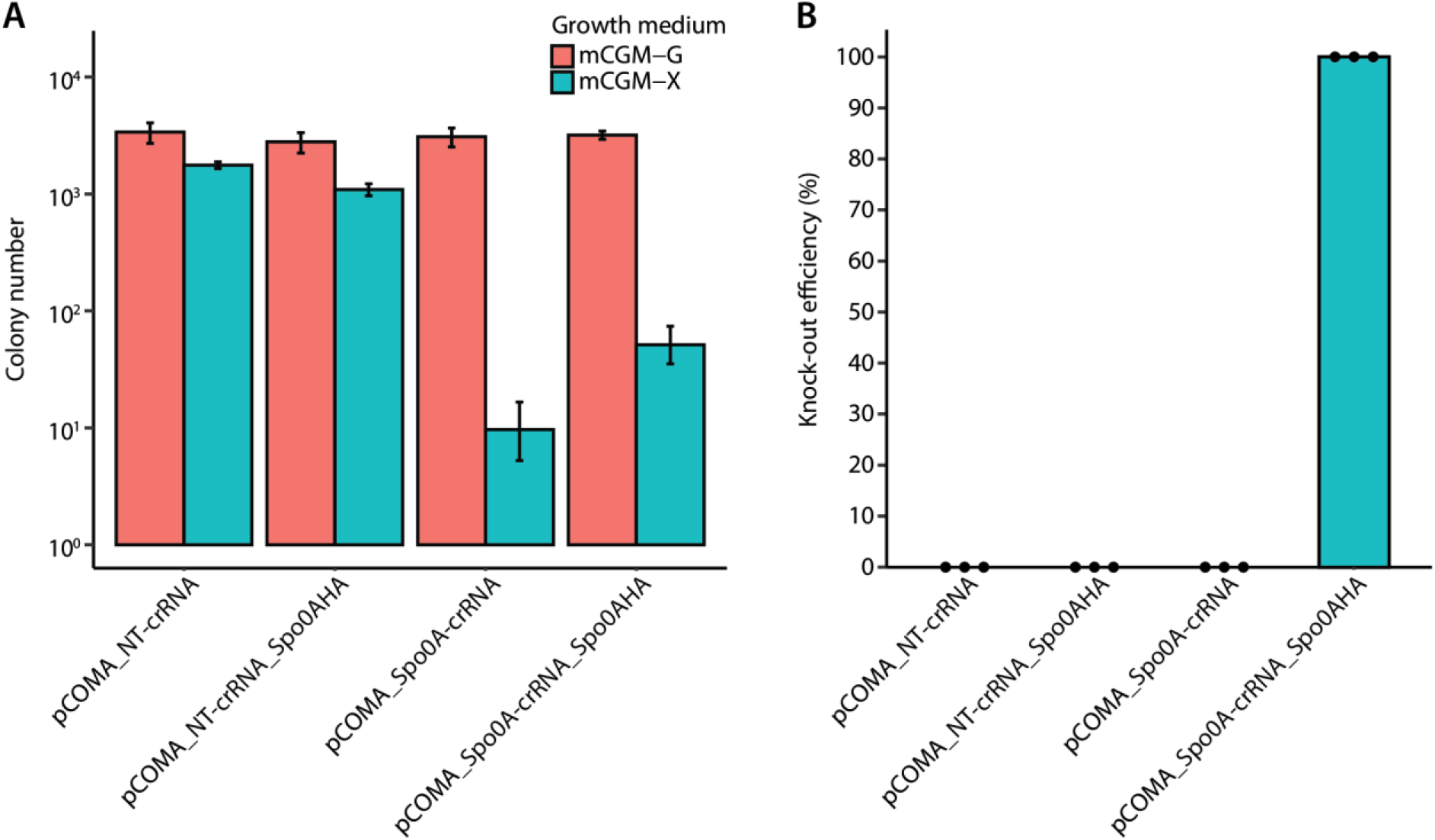
Inducible counterselection and *Spo0A* knockout in *C. beijerinckii* NCIMB 8052. *C. beijerinckii* NCIMB 8052 cells were transformed either with the pCOMA_NT-crRNA and pCOMA_NT-crRNA_Spo0AHA non-targeting plasmids (negative controls), the pCOMA_Spo0A-crRNA targeting plasmid (positive control) or the pCOMA_Spo0A-crRNA_Spo0AHA plasmid. Transformants were plated on mCGM-G (no-induction) or on mCGM-X (induction of FnCas12a). This experiment was performed in biological triplicates. The error bars in (A) show the standard deviation. (**A**) Colony number obtained after plating the transformants on the appropriate medium. (**B**) *Spo0A* knockout efficiency determined by screening eight colonies from each replicate. Dots represent the knockout efficiency from each replicate.

To assess whether the colonies obtained on mCGM-X were successful *Spo0A* knockouts, we screened eight colonies (if present) from each biological replicate (24 in total) through colony PCR. As expected, the non-targeting controls showed a 0% knockout efficiency represented by a WT genotype of 1866 bp amplicons (Fig. 2B and S2). Similarly, the obtained pCOMA_Spo0A-crRNA colonies showed a WT genotype (0% knockout efficiency), further supporting that the obtained colonies are escapees. Intriguingly, obtained colonies containing the pCOMA_Spo0A-crRNA_Spo0AHA had a 100% knockout efficiency with a Δ*Spo0A* genotype corresponding to 1044 bp amplicons (Fig. 2B and S2). Mutant colonies were further assessed for the deletion of *Spo0A* by Sanger sequencing, confirming the scarless deletion of *Spo0A*.

### Δ*Spo0a C. beijerinckii* NCIMB 8052 shows retarded growth, elimination of solvent production and increased production of acids

As previously described (32,33), Δ*Spo0A C. beijerinckii* strains show a distinctive phenotype which includes the elimination of solvent production, increased acid production and altered colony morphology. To assess whether our Δ*Spo0A C. beijerinckii* NCIMB 8052 mutants show the described phenotypic characteristics, we selected three Δ*Spo0A* colonies and subjected them to plasmid curing. Three cured Δ*Spo0A C. beijerinckii* NCIMB 8052 colonies and three WT *C. beijerinckii* NCIMB 8052 colonies were then grown in GAPES medium for 48 h and the fermentation products were analysed (Fig. 3A and B).

**Figure 3.**
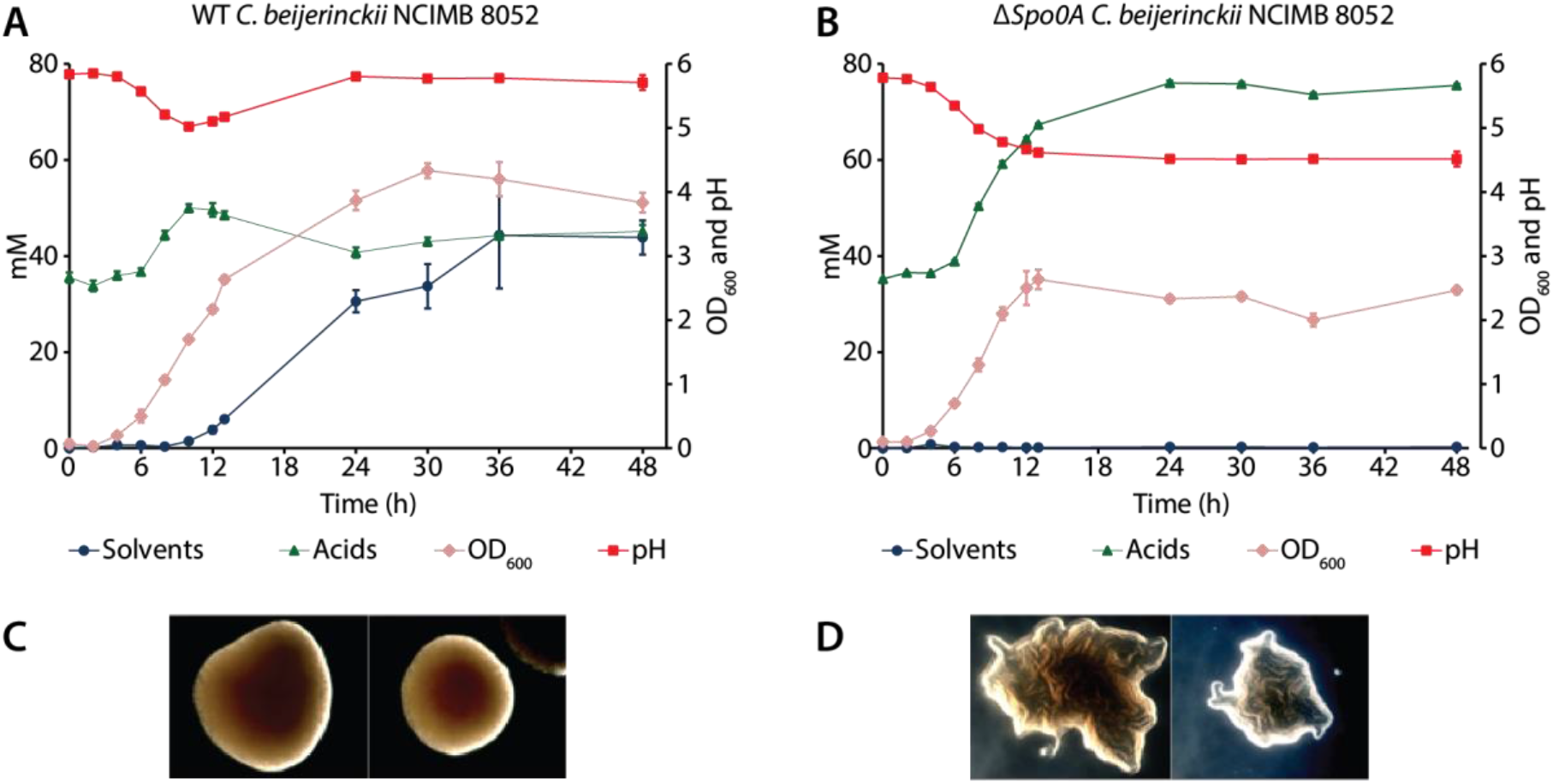
Fermentation profile and morphology of WT and Δ*Spo0A C. beijerinckii* NCIMB 8052 strains. **(A)** Fermentation profile of WT *C. beijerinckii* NCIMB 8052. (**B**) Fermentation profile of Δ*Spo0A C. beijerinckii* NCIMB 8052. The error bars in (A) and (B) indicate the standard deviation. Solvents represent acetone, butanol and ethanol. Acids represent acetate, butyrate and lactate. (**C**) Colony morphology of WT *C. beijerinckii* NCIMB 8052. (**D**) Colony morphology of Δ*Spo0A C. beijerinckii* NCIMB 8052.

During the first 8 h of incubation, the Δ*Spo0A* and WT strains performed similarly, as the acids were produced to equimolar amounts (approx. 50 mM), accompanied with the characteristic pH drop from pH 6.0 to around pH 5.0. However, after 8 h of incubation, the growth and product formation of the Δ*Spo0A* and WT strains differed considerably. The WT strain ceased the production of acids and the production of solvents was initiated, raising the pH to around 5.8. The OD_600_ increased to 4.33 at 30 h of incubation and it decreased to 3.83 at the end of the fermentation. Solvents and acids reached a concentration of 43.85 mM and 45.11 mM, respectively, at the end of fermentation. In contrast, the Δ*Spo0A* strain did not assimilate the produced acids after 8 h of growth, resulting in an increased production of acids and the absence of solvent production. Due to the lack of acid assimilation, the pH remained low, reaching a pH of 4.51 at the end of the fermentation. The OD_600_ did not increase after 12 h of fermentation, likely due to the acid crash caused by the increased acid production. The acids reached a concentration of 75.5 mM at the end of the fermentation and no solvents were produced. Lastly, the distinct, snowflake-like colony phenotype was apparent for the Δ*Spo0A* colonies whereas the WT colonies showed the typical round and smooth shape (Fig. 3C and D).

### Establishing single deletions of various *C. beijerinckii* NCIMB 8052 genes

To assess the applicability and knockout efficiency of our tool to other genes (other than the *Spo0A*), we sought to delete four genes at different genomic loci: *Cbei_0408* (*Upp*; 630 bp), *Cbei_1291* (987 bp), *Cbei_3238* (888 bp) and *Cbei_3932* (813 bp). An identical protocol as the one described for *Spo0A* deletion was followed and obtained colonies were screened for mutants through colony PCR and Sanger sequencing.

In contrast to the *Spo0A* knockouts, the knockout efficiency varied amongst the selected genes. The highest knockout efficiency was observed for *Upp* (79%), followed by *Cbei_3238* (42%), *Cbei_3932* (38%) and *Cbei_1291* (25%) (Fig. 4 and S3). More intriguingly, we clearly observed different knockout efficiencies between the biological replicates of transformants carrying (essentially) the same plasmid variant. For example, for *Cbei_1291*, two out of the three biological replicates showed 0% knockout efficiency, whereas one of the replicates showed 75% knockout efficiency (Fig. S3). The inconsistency in editing amongst the replicates was also observed for *Cbei_3238* and *Cbei_3932* where at least one of the replicates showed 0% knockout efficiency. Nonetheless, one of the replicates for *Cbei_3238* showed 88% knockout efficiency and one of the replicates for *Cbei_3932* showed 63% knockout efficiency. Knocking out the *Upp* gene was more consistent as biological replicates varied between 63 and 100% knockout efficiency.

**Figure 4.**
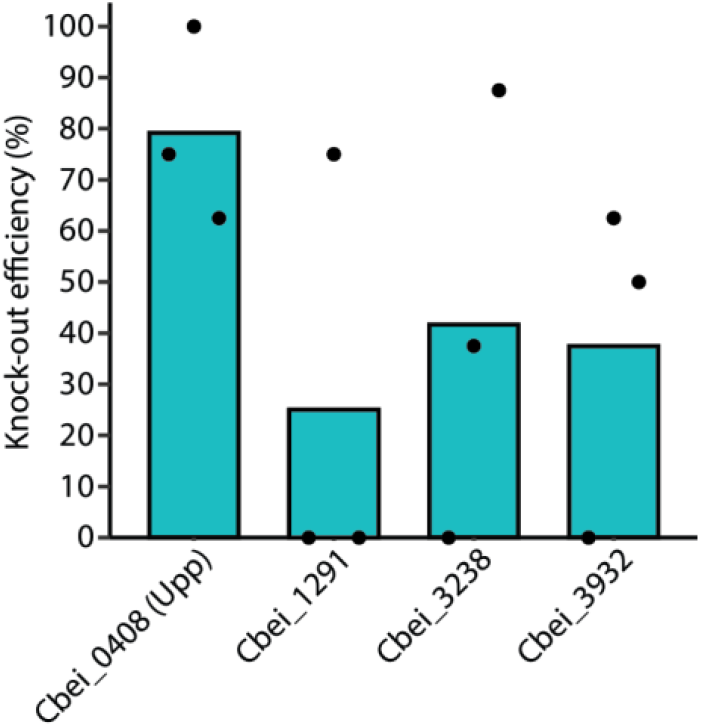
Single-gene knockout of multiple genes in *C. beijerinckii* NCIMB 8052. *Cbei_0408* (*Upp*), *Cbei_1291, Cbei_3238* and *Cbei_3932* were targeted for knockout. The average knockout efficiency for each gene is: *Cbei_0408* (79.17%), *Cbei_1291* (25%), *Cbei_3238* (41.67%) and *Cbei_3932* (37.5%). This experiment was performed in biological triplicates. The knockout efficiency was determined by screening eight colonies from each replicate through colony PCR using the primers listed in Table S1. Dots represent the knockout efficiency from each replicate.

The variable knockout efficiency amongst the different genes may be attributed to the selection of a good or bad spacer (34) which is often attributed to the secondary structure of the crRNA, the GC content of the protospacer and the spacer, and the melting temperature of the spacer-protospacer pairing (35,36). However, the variable knockout efficiency amongst the biological replicates targeting the same gene could be due to early escapees which dominated the culture during the 48 h growth before inducing the expression of FnCas12a for counterselection. Unfortunately, we did not analyse the cause of escapees but potential reasons may include mutations at the spacer or protospacer, at the PAM sequence or the *FnCas12* gene sequence.

### Multiplex gene knockout in *C. beijerinckii* NCIMB 8052

Single-gene knockout in *C. beijerinckii* NCIMB 8052 was previously achieved using CRISPR-AsCas12a (15). However, multiplex genome editing was never demonstrated before. To show that multiplex gene knockout is possible in *C. beijerinckii* NCIMB 8052, we sought to knockout the *Spo0A* and *Upp* genes in a single step (i.e., in one transformation event) using our established xylose-inducible system as described above.

To target two genomic sequences (*Spo0A* and *Upp*) with FnCas12a, two spacers (one for each target) were introduced into the CRISPR array (Table S1). Since the spacer sequence can affect the editing efficiency (37), we constructed a CRISPR array where the Spo0A spacer preceded the Upp1 spacer (Spo0A-Upp1) and a CRISPR array where the Upp spacer preceded the Spo0A spacer (Upp1-Spo0A). In addition, to further assess the effect of changing one of the targeting spacers with another spacer that targets the same gene but in a different genomic location, we replaced the Upp1 spacer with the Upp2 spacer. Similarly, a CRISPR array where the Spo0A spacer preceded the Upp2 spacer (Spo0A-Upp2) and a CRISPR array where the Upp2 spacer preceded the Spo0A (Upp2-Spo0A) were constructed (Table S1).

As expected, knockout efficiencies varied between the different CRISPR array variants (Fig. 5). The highest knockout efficiency for *Upp* (91%) was observed when the Spo0A-Upp1 CRISPR array was used, whereas the lowest knockout efficiency (13%) was observed when the Spo0A-Upp2 CRISPR array was used. While the Spo0A-Upp1 CRISPR array showed the highest knockout efficiency for *Upp*, 0% knockout efficiency was observed for *Spo0A*. In contrast, by switching the position of the Spo0A and Upp1 spacers (i.e., from Spo0A-Upp1 to Upp1-Spo0A), 18% knockout efficiency was observed for *Spo0A*. The knockout efficiency of *Upp* dropped to 77% when the Upp1-Spo0A CRISPR array was used, indicating that the position of the Upp1 spacer in the array does not affect largely the knockout efficiency of *Upp*. Following, the knockout efficiency of *Upp* was reduced from 92% to 13% when the Upp1 spacer was substituted with the Upp2 spacer and when the Spo0A spacer was the first spacer of the CRISPR array. However, the substitution of Upp1 to Upp2 yielded some (5/24) successful Spo0A knockouts, although most (4/5) of them were mixed colonies, as indicated in Fig. S4. Important to note is that half (12/24) of the screened colonies had a mixed (WT and Δ*Upp*) genotype.

**Figure 5.**
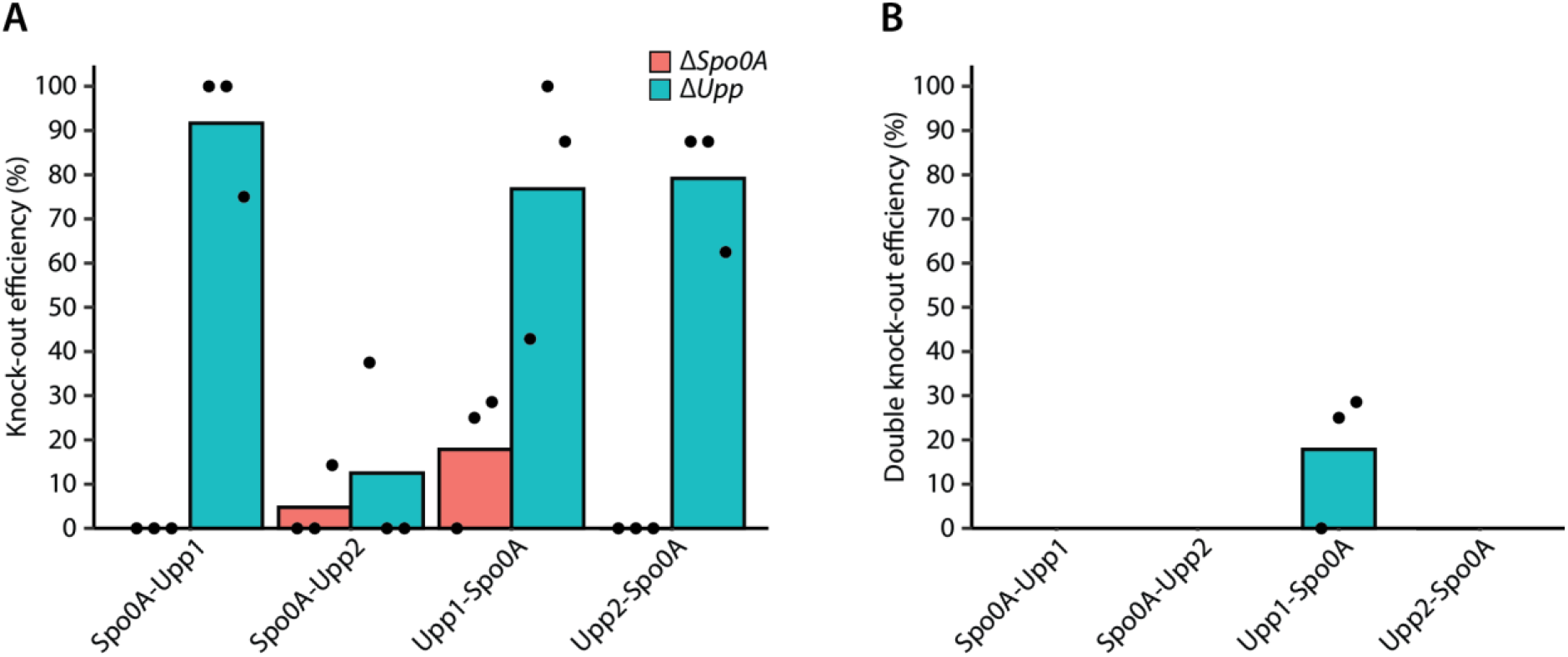
Multiplex gene knockout. The *Spo0A* and *Upp* genes were targeted simultaneously for knockout in a single step. Different CRISPR arrays were used with either the Spo0A spacer preceding the Upp spacer or the other way around. Two different Upp spacers were used, designated as Upp1 and Upp2. (**A**) Knockout efficiency of either the *Spo0A* or the *Upp* gene using the different CRISPR arrays. (**B**) Double knockout efficiency of the *Spo0A* and *Upp* genes using the different CRISPR arrays. The knockout efficiency was determined by screening eight colonies from each replicate through colony PCR using the primers listed in Table S1. Dots represent the knockout efficiency from each replicate.

In summary, we achieved multiplex gene knockout in *C. beijerinckii* NCIMB 8052, although with low editing efficiency. Clean (i.e., without the presence of mixed colonies) double *Spo0A* and *Upp* mutants were observed only when the Upp1-Spo0A CRISPR array was used. This observation is not surprising as a previous report by Liao et al. (2019) clearly demonstrated that the abundance of crRNAs in a CRISPR array varies widely. The variation in the abundance of crRNAs is very likely to be due to the effect of secondary structures which inhibit the formation of the characteristic hairpin required for Cas12a processing (18,37). To assess this possibility, we predicted the secondary structure of the transcribed pre-crRNAs using NUPACK (38).

As expected, complex secondary structures were formed in all the pre-crRNAs (Fig. S5). In the CRISPR arrays where the Spo0A spacer preceded the Upp spacer, an undisrupted hairpin was formed between nucleotide 22 and 35 of the pre-crRNA, recommending the successful recognition and processing of the Spo0A crRNA by FnCas12a. However, a secondary structure was observed between the nucleotides of the Spo0A spacer, although with low equilibrium probability (Fig. S5). The secondary structure formed by the Spo0A spacer may reflect the low knockout efficiency observed in all the multiplex editing assays, rendering the Spo0A crRNA as a poorly performing crRNA (34). Yet, 100% knockout efficiency was observed when the Spo0A crRNA was used for single *Spo0A* knockouts (Fig. 2B). In contrast to the Spo0A spacer, the necessary hairpin for processing the Upp1 or Upp2 crRNA was disturbed in the Spo0A-Upp1 and Spo0A-Upp2 arrays. Still, our results for both the Spo0A-Upp1 and Spo0A-Upp2 arrays show that the (hypothetically) well processed Spo0A crRNA yields low knockout efficiency for the *Spo0A* gene, whereas the disturbed Upp1 or Upp2 crRNAs are not necessarily a limitation for knocking out the *Upp* gene (Fig. 4A). When the Upp2 spacer preceded the Spo0A spacer, disturbed hairpins were observed for both the Upp2 and Spo0A crRNAs (Fig. S5). In contrast, in the case where the Upp1 spacer preceded the Spo0A spacer, undisturbed hairpins were formed for both crRNAs. The Upp1-Spo0A array combination yielded the highest (18%) multiplex knockout efficiency (Fig. 5B), which is very likely to be the result of undisturbed hairpins.

In total, our multiplex knockout results cannot be fully explained by the pre-crRNA secondary structure. In most cases, the Spo0A crRNA hairpin is structured but yields very low knockout efficiency, whereas the Upp crRNA hairpin is unstructured and often yields high knockout efficiency. Based on our observations and the observations made by Liao et al. (2019) and Creutzburg et al. (2020), we can conclude that when a multiplex approach is considered, the test of multiple spacers targeting the same gene in combination with the change in spacer position in the CRISPR array should be applied for optimal results.

## Conclusion

In this study, we successfully developed a CRISPR-FnCas12a genome engineering tool for *C. beijerinckii* NCIMB 8052 that can facilitate single- and multi-plex gene knockout in a single step. The knockout efficiency for single genes varied between 25 and 100%, indicating that different genomic loci are not targeted and deleted equally. The knockout efficiency for the simultaneous deletion of two genes was 18% and dependentent on the spacer sequence and position. In general, our tool expands the CRISPR-Cas toolbox in *Clostridia* species and can contribute to the rapid and easy generation of mutants.

## Acknowledgements

We would like to thank Rob Joosten and Ton van Gelder for their technical support on this project. Also, we would like to thank Dr. Wen Wu and Dr. Prarthana Mohanraju for their advice and for sharing their knowledge on CRISPR-Cas12a.

## Supplementary information

**Supplementary Figure 1.**
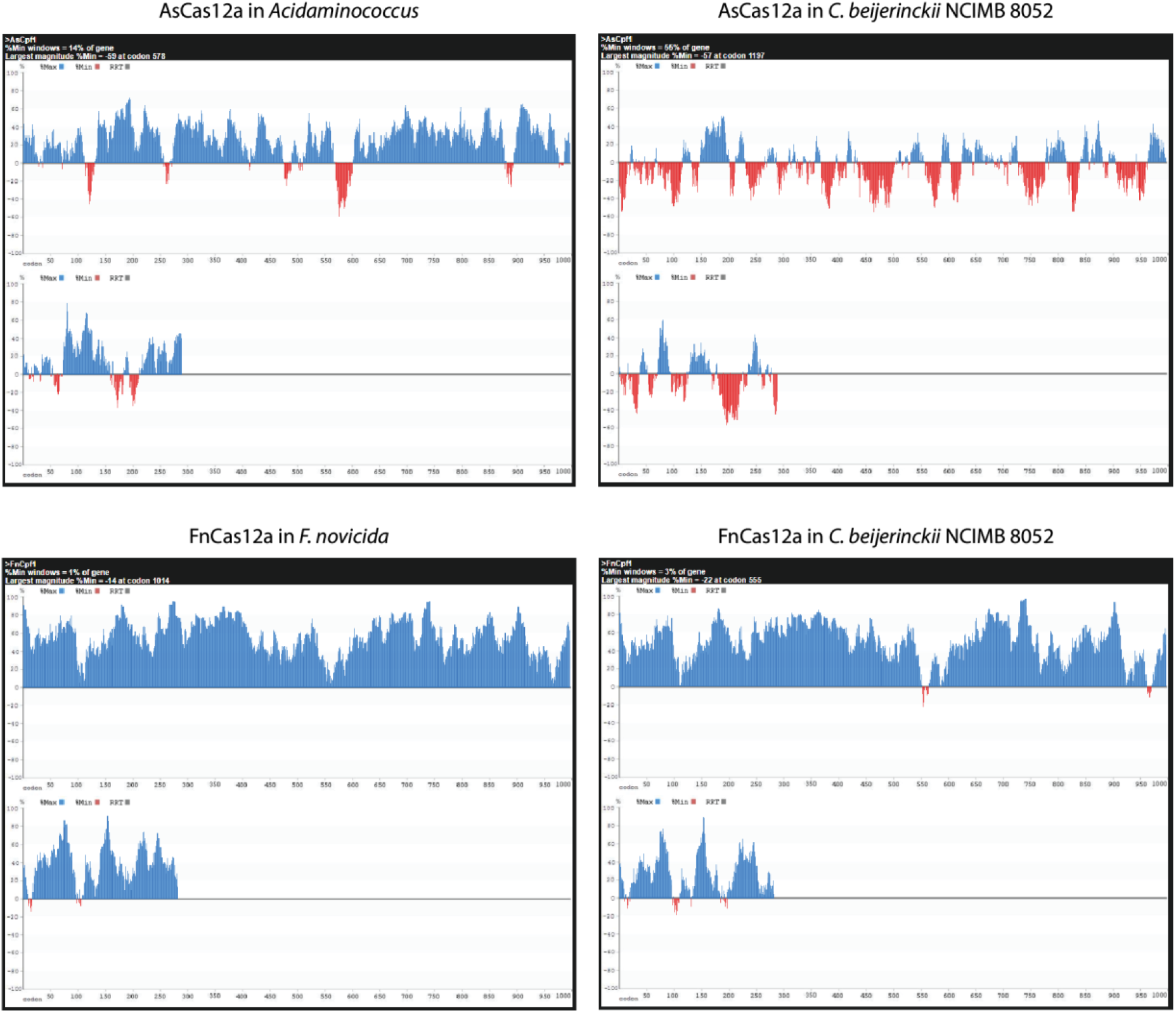
Codon usage of AsCas12a and FnCas12a in the native (*Acidaminococcus* or *Francisella novicida*) and target (*C. beijerinckii* NCIMB 8052) organisms. The codon usage for each organism was created using the codon harmonization tool developed by Claassens et al. (2017). The Cas12a codon usage in the native and target organisms was visualized using the codons.org website created by Clarke T.F. & Clark P.L. (2008).

**Supplementary Figure 2.**
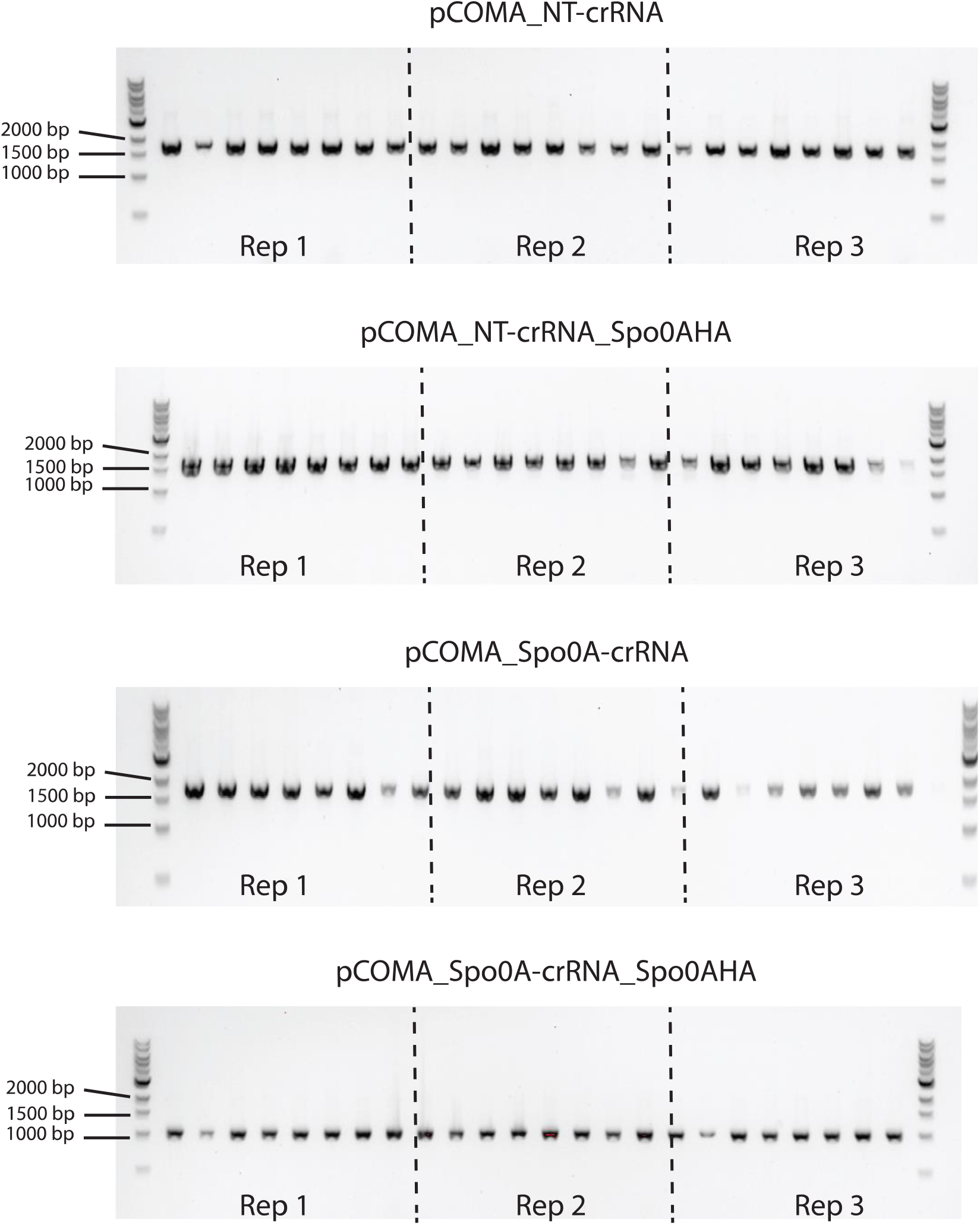
Raw data for the single-gene knockout of *Spo0A* using CRISPR-FnCas12a in *C. beijerinckii* NCIMB 8052. *C. beijerinckii* NCIMB 8052 was transformed either with pCOMA_NT-crRNA, pCOMA_NT-crRNA_Spo0AHA, pCOMA_Spo0A-crRNA or pCOMA_Spo0A-crRNA_Spo0AHA and obtained colonies were screened through colony PCR using BG16483 and BG16484 oligos (Table S1). This experiment was performed in biological triplicates and the result of each triplicate is represented by Rep1, Rep2 and Rep3 at the bottom of each gel and separated by the dashed vertical lines. Mix amplicons (wild-type and knockout bands) were not counted for the total knockout efficiency percentage. Wild-type Spo0A: 1866 bp, ΔSpo0A: 1044 bp.

**Supplementary Figure 3.**
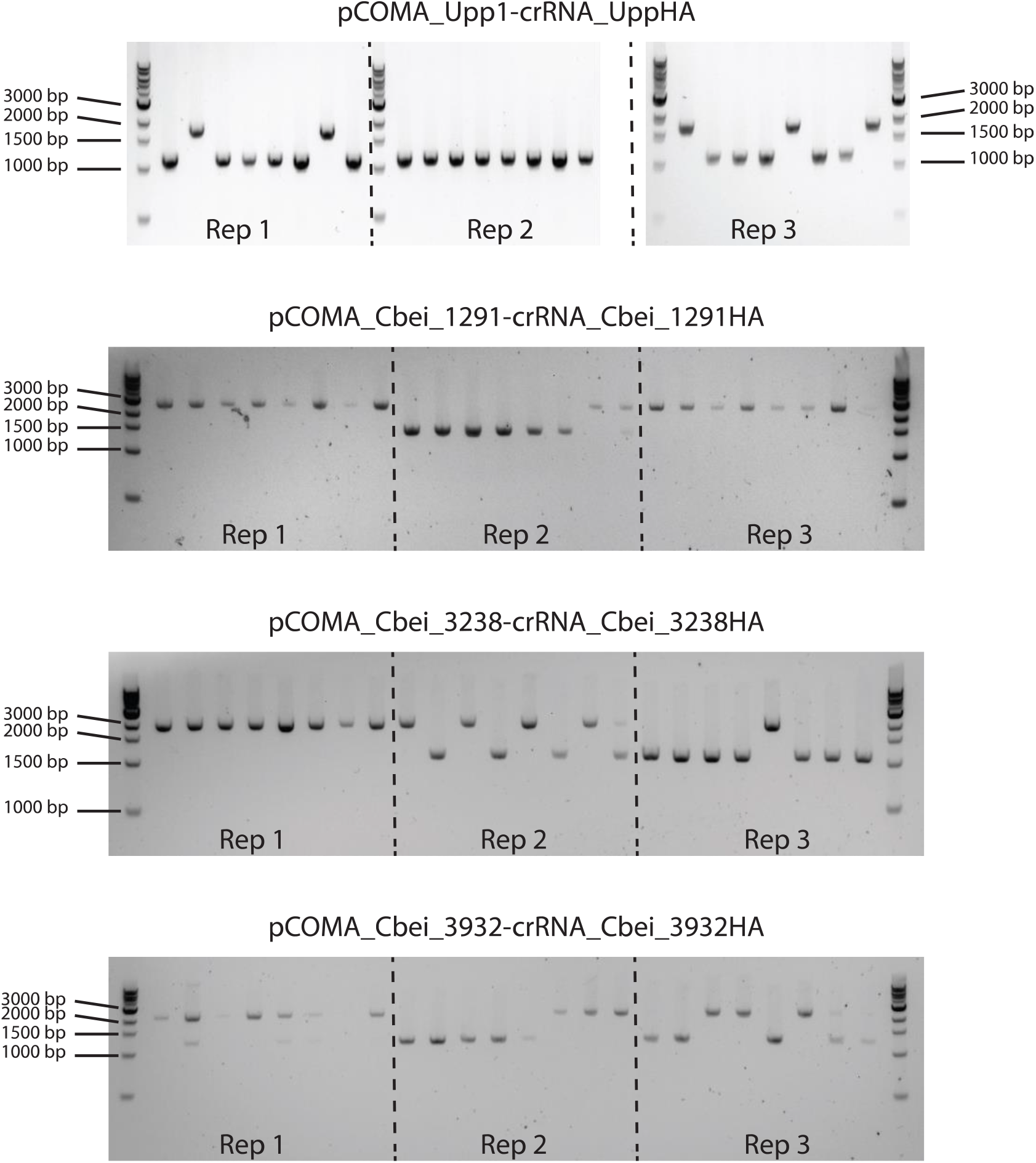
Raw data for the single-gene knockout of the *Upp, Cbei_1291, Cbei_3238* and *Cbei_3932* genes using CRISPR-FnCas12a in *C. beijerinckii* NCIMB 8052. *C. beijerinckii* NCIMB 8052 was transformed with either of pCOMA_Upp1-crRNA_UppHA, pCOMA_Cbei_1291-crRNA_Cbei_1291HA, pCOMA_Cbei_3238-crRNA_ Cbei_3238HA or pCOMA_Cbei_3932-crRNA_Cbei_3932HA. Each knockout experiment was performed in biological triplicates and the result of each triplicate is represented by Rep1, Rep2 and Rep3 at the bottom of each gel and separated by the dashed vertical lines. Mix amplicons (wild-type and knockout bands) were not counted for the total knockout efficiency percentage. The primers used for this experiment are listed in Table S1. Wild-type *Upp*: 1775 bp, Δ*Upp*: 1145 bp. Wild-type *Cbei_1291*: 2392 bp, Δ*Cbei_1291*: 1405 bp. Wild-type *Cbei_3238*: 1979 bp, Δ*Cbei_3238*: 1091 bp. Wild-type *Cbei_3932*: 2054 bp, Δ*Cbei_3932*: 1241 bp.

**Supplementary Figure 4.**
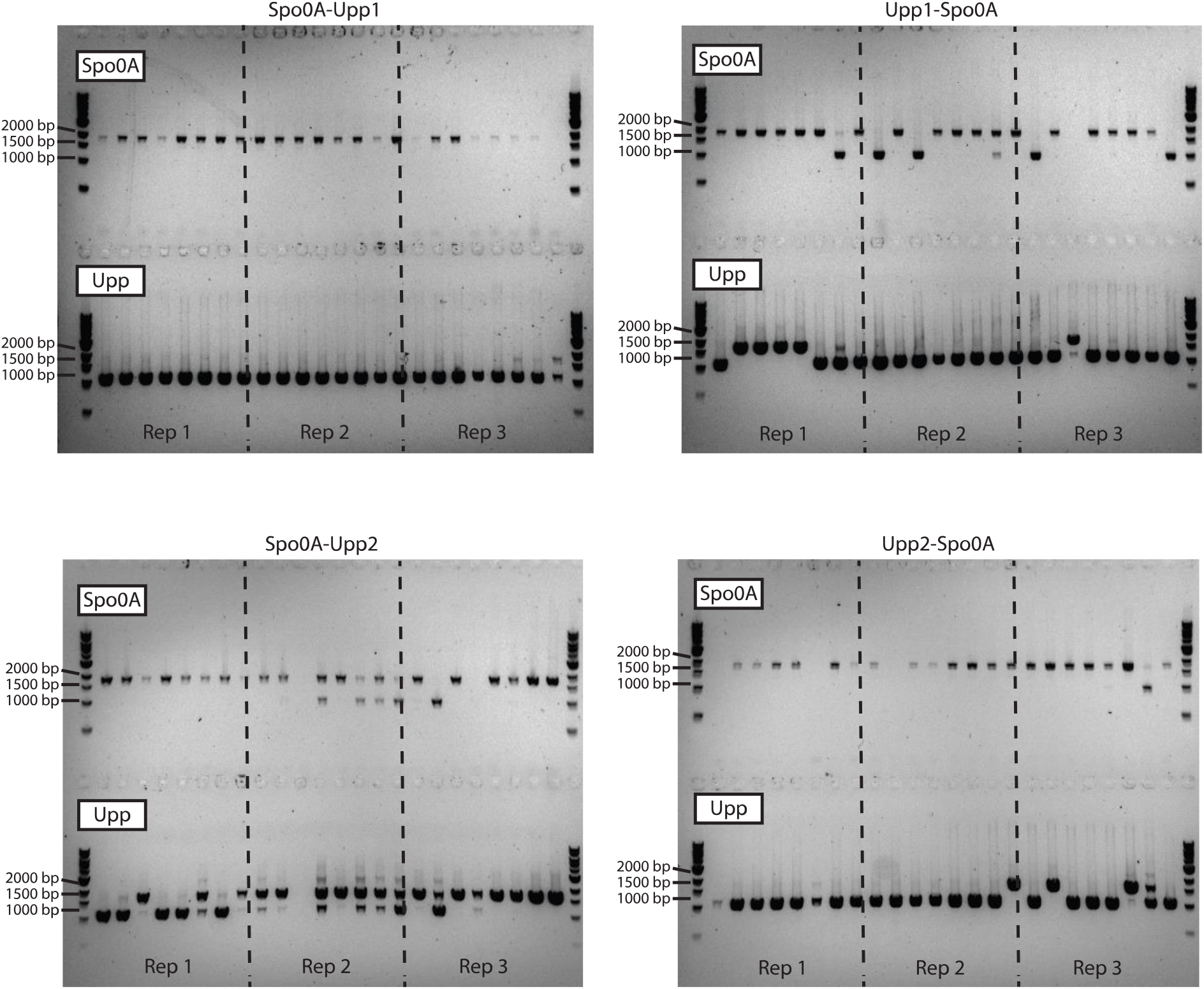
Raw data for the multiplex knockout of *Spo0A* and *Upp* using CRISPR-FnCas12a in *C. beijerinckii* NCIMB 8052. pCOMA_Spo0A-Upp1-crRNA_Spo0AHA_UppHA (top left), pCOMA_Upp1-Spo0A-crRNA_Spo0AHA_UppHA (top right), pCOMA_Spo0A-Upp2-crRNA_Spo0AHA_UppHA (bottom left) or pCOMA_Upp2-Spo0A-crRNA_Spo0AHA_UppHA (bottom right) was used to transform *C. beijerinckii* NCIMB 8052. Each knockout experiment was performed in biological triplicates and the result of each triplicate is represented by Rep1, Rep2 and Rep3 at the bottom of each gel and separated by the dashed vertical lines. Mix amplicons (wild-type and knockout bands) were not counted for the total knockout efficiency percentage. The primers used for this experiment are listed in Table S1. Wild-type *Spo0A*: 1866 bp, Δ*Spo0A*: 1044 bp. Wild-type *Upp*: 1775 bp, Δ*Upp*: 1145 bp.

**Supplementary Figure 5.**
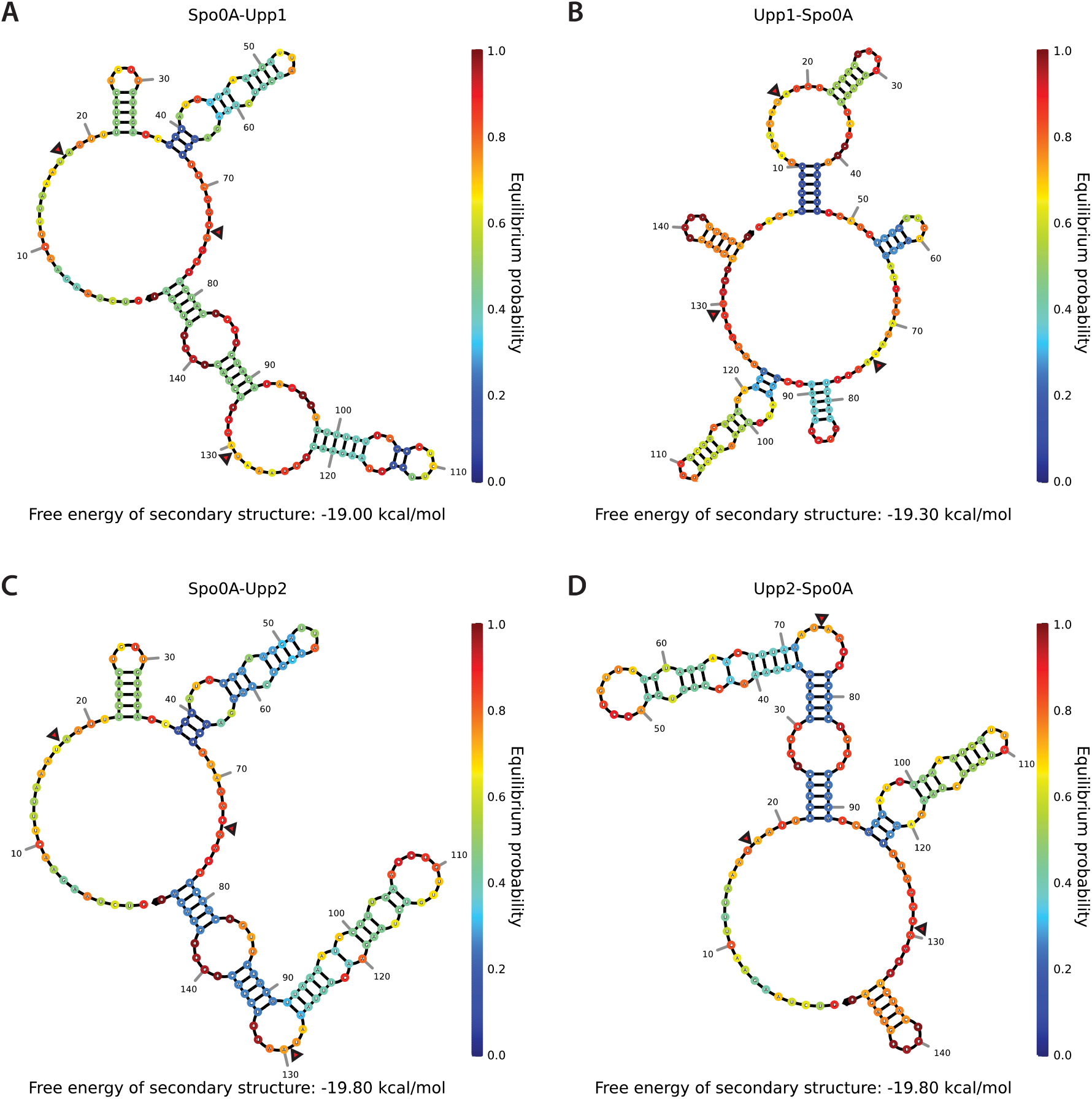
Predicted secondary structure of pre-crRNAs. The pre-crRNAs include the three 36 nt repeats interspaced with the designated 20 nt spacers. Red triangles indicate the processing site by FnCas12a. A) pre-crRNA where the Spo0A spacer precedes the Upp1 spacer. B) pre-crRNA where the Upp1 spacer precedes the Spo0A spacer. C) pre-crRNA where the Spo0A spacer precedes the Upp2 spacer. D) pre-crRNA where the Upp2 spacer precedes the Spo0A spacer.

**Supplementary Table 1.**
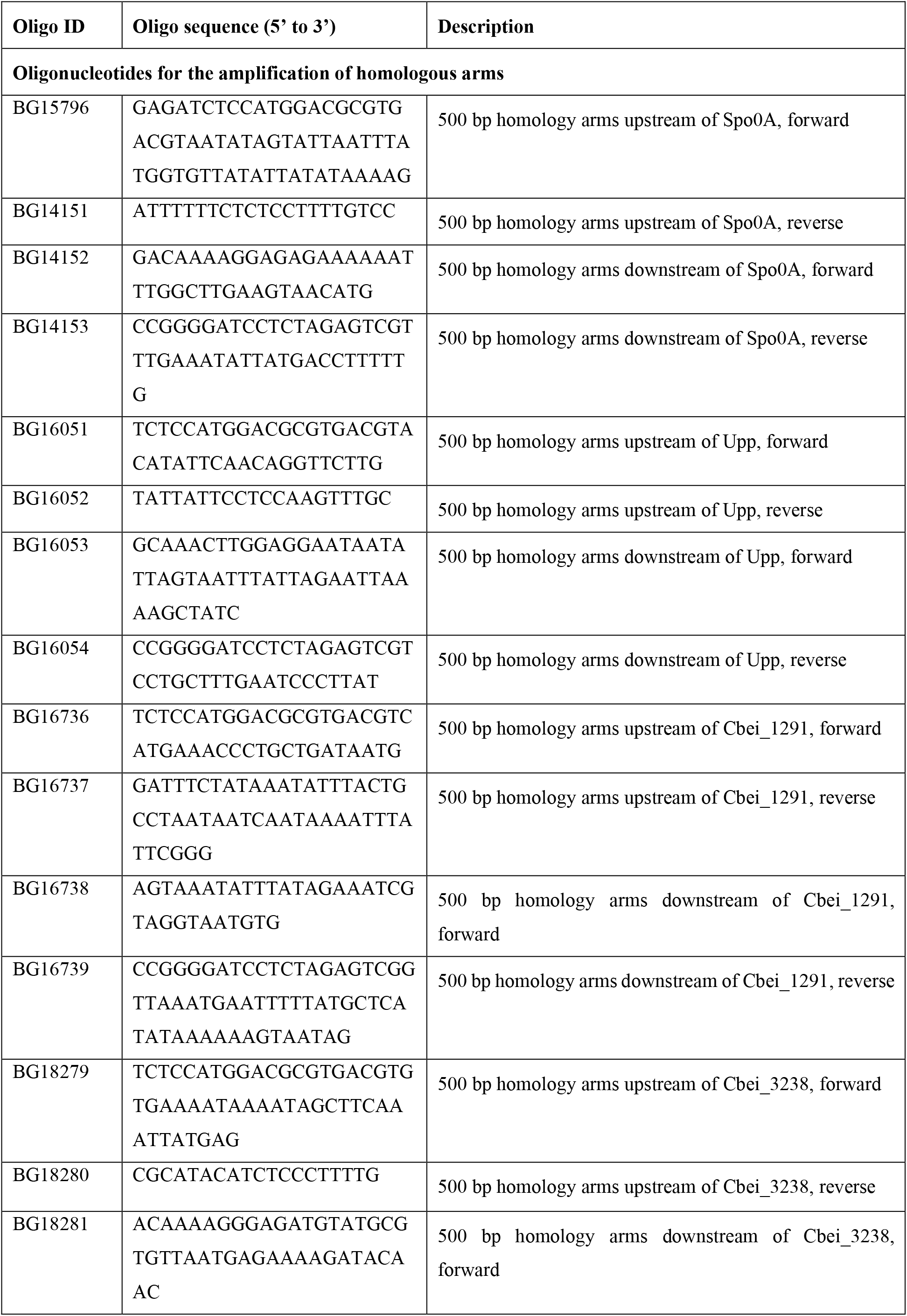
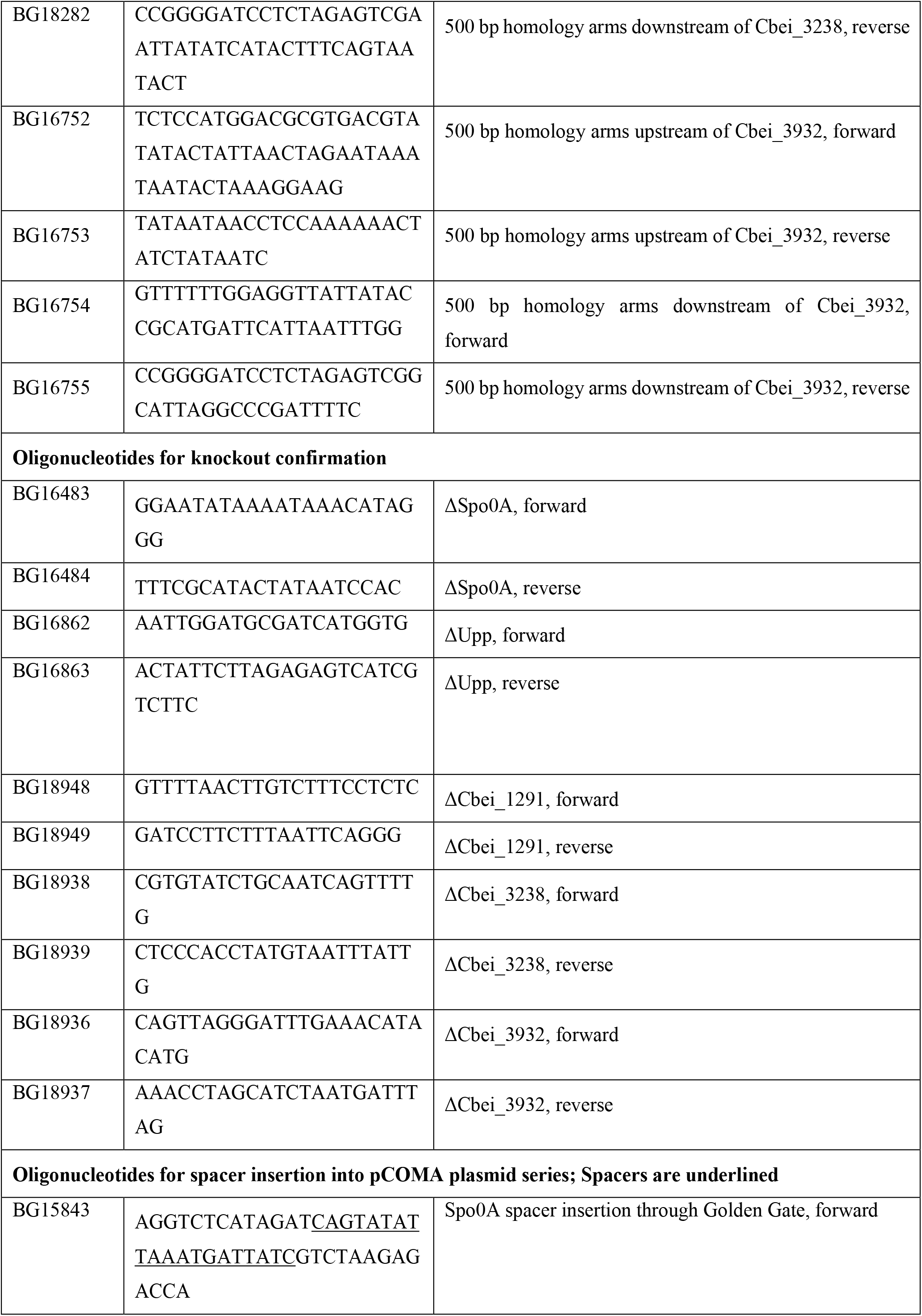
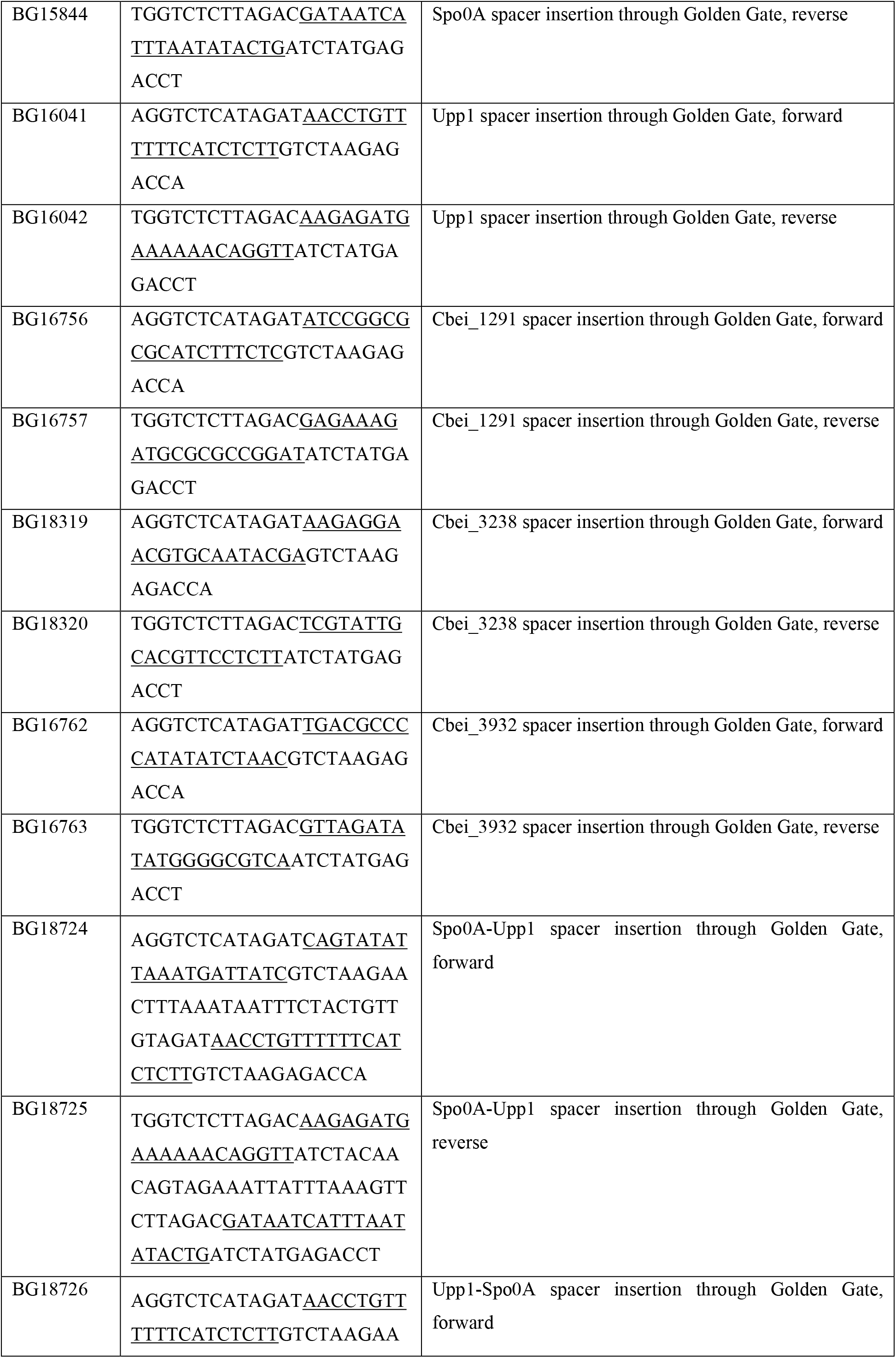
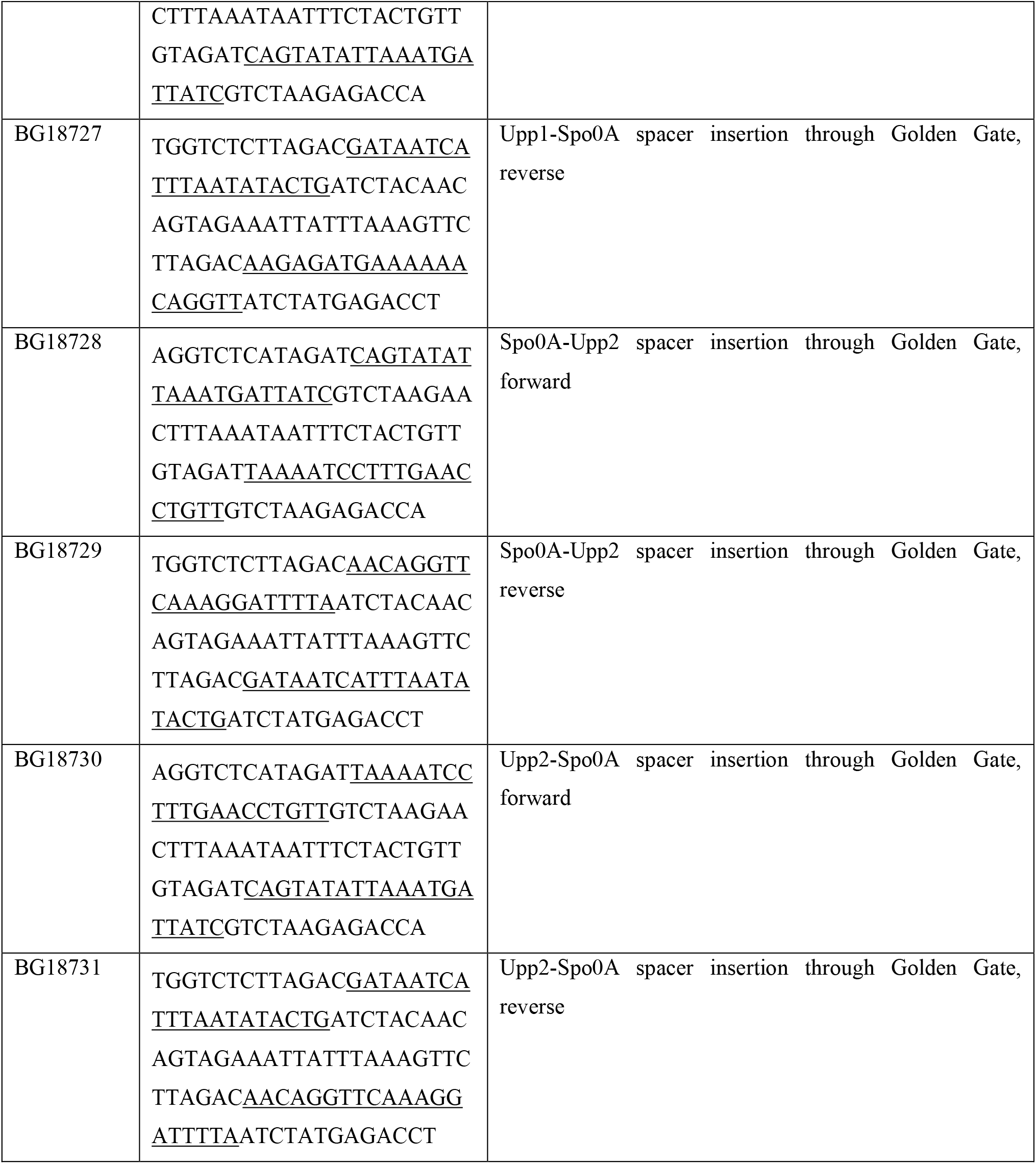
Oligonucleotides used in this study.

## References

1. Gabriel, C. (1928) Butanol fermentation process1. Industrial & engineering chemistry, 20, 1063–1067.

2. Gibbs, D. (1983) The rise and fall (… and rise?) of acetone/butanol fermentations. Trends in Biotechnology, 1, 12–15.

3. Killeffer, D. (1927) Butanol and acetone from corn1: a description of the fermentation process. Industrial & Engineering Chemistry, 19, 46–50.

4. Spivey, M. (1978) Acetone/butanol/ethanol fermentation. Process Biochem.;(United Kingdom), 13.

5. Nolling, J., Breton, G., Omelchenko, M.V., Makarova, K.S., Zeng, Q., Gibson, R., Lee, H.M., Dubois, J., Qiu, D. and Hitti, J. (2001) Genome sequence and comparative analysis of the solvent-producing bacterium Clostridium acetobutylicum. Journal of bacteriology, 183, 4823–4838.

6. McAllister, K.N. and Sorg, J.A. (2019) CRISPR genome editing systems in the genus Clostridium: a timely advancement. Journal of bacteriology, 201, e00219–00219.

7. Heap, J.T., Pennington, O.J., Cartman, S.T., Carter, G.P. and Minton, N.P. (2007) The ClosTron: a universal gene knock-out system for the genus Clostridium. Journal of microbiological methods, 70, 452–464.

8. Heap, J.T., Cartman, S.T., Kuehne, S.A., Cooksley, C. and Minton, N.P. (2010), Clostridium difficile. Springer, pp. 165–182.

9. Wang, Y., Zhang, Z.-T., Seo, S.-O., Choi, K., Lu, T., Jin, Y.-S. and Blaschek, H.P. (2015) Markerless chromosomal gene deletion in Clostridium beijerinckii using CRISPR/Cas9 system. Journal of biotechnology, 200, 1–5.

10. Jiang, W., Bikard, D., Cox, D., Zhang, F. and Marraffini, L.A. (2013) RNA-guided editing of bacterial genomes using CRISPR-Cas systems. Nature biotechnology, 31, 233–239.

11. Li, Q., Chen, J., Minton, N.P., Zhang, Y., Wen, Z., Liu, J., Yang, H., Zeng, Z., Ren, X. and Yang, J. (2016) CRISPR-based genome editing and expression control systems in Clostridium acetobutylicum and Clostridium beijerinckii. Biotechnology journal, 11, 961–972.

12. Wang, Y., Zhang, Z.-T., Seo, S.-O., Lynn, P., Lu, T., Jin, Y.-S. and Blaschek, H.P. (2016) Bacterial genome editing with CRISPR-Cas9: deletion, integration, single nucleotide modification, and desirable “clean” mutant selection in Clostridium beijerinckii as an example. ACS synthetic biology, 5, 721–732.

13. Li, Q., Seys, F.M., Minton, N.P., Yang, J., Jiang, Y., Jiang, W. and Yang, S. (2019) CRISPR–Cas9D10A nickase-assisted base editing in the solvent producer Clostridium beijerinckii. Biotechnology and bioengineering, 116, 1475–1483.

14. Wang, Y., Zhang, Z.T., Seo, S.O., Lynn, P., Lu, T., Jin, Y.S. and Blaschek, H.P. (2016) Gene transcription repression in Clostridium beijerinckii using CRISPR-dCas9. Biotechnology and bioengineering, 113, 2739–2743.

15. Zhang, J., Hong, W., Zong, W., Wang, P. and Wang, Y. (2018) Markerless genome editing in Clostridium beijerinckii using the CRISPR-Cpf1 system. Journal of biotechnology, 284, 27–30.

16. Diallo, M., Hocq, R., Collas, F., Chartier, G., Wasels, F., Wijaya, H.S., Werten, M.W., Wolbert, E.J., Kengen, S.W. and van der Oost, J. (2020) Adaptation and application of a two-plasmid inducible CRISPR-Cas9 system in Clostridium beijerinckii. Methods, 172, 51–60.

17. Jinek, M., Chylinski, K., Fonfara, I., Hauer, M., Doudna, J.A. and Charpentier, E. (2012) A programmable dual-RNA–guided DNA endonuclease in adaptive bacterial immunity. science, 337, 816–821.

18. Zetsche, B., Gootenberg, J.S., Abudayyeh, O.O., Slaymaker, I.M., Makarova, K.S., Essletzbichler, P., Volz, S.E., Joung, J., Van Der Oost, J. and Regev, A. (2015) Cpf1 is a single RNA-guided endonuclease of a class 2 CRISPR-Cas system. Cell, 163, 759–771.

19. Gapes, J.R., Nimcevic, D. and Friedl, A. (1996) Long-Term Continuous Cultivation of Clostridium beijerinckii in a Two-Stage Chemostat with On-Line Solvent Removal. Appl. Environ. Microbiol., 62, 3210–3219.

20. Scotcher, M.C. and Bennett, G.N. (2005) SpoIIE regulates sporulation but does not directly affect solventogenesis in Clostridium acetobutylicum ATCC 824. Journal of bacteriology, 187, 1930–1936.

21. Batianis, C., Kozaeva, E., Damalas, S.G., Martín-Pascual, M., Volke, D.C., Nikel, P.I. and Martins dos Santos, V.A. (2020) An expanded CRISPRi toolbox for tunable control of gene expression in Pseudomonas putida. Microbial Biotechnology, 13, 368–385.

22. Oultram, J., Loughlin, M., Swinfield, T., Brehm, J., Thompson, D. and Minton, N. (1988) Introduction of plasmids into whole cells of Clostridium acetobutylicum by electroporation. FEMS microbiology letters, 56, 83–88.

23. Diallo, M., Hocq, R., Collas, F., Chartier, G., Wasels, F., Wijaya, H.S., Werten, M.W., Wolbert, E.J., Kengen, S.W. and van der Oost, J. (2019) Adaptation and application of a two-plasmid inducible CRISPR-Cas9 system in Clostridium beijerinckii. Methods.

24. Claassens, N.J., Siliakus, M.F., Spaans, S.K., Creutzburg, S.C., Nijsse, B., Schaap, P.J., Quax, T.E. and Van Der Oost, J. (2017) Improving heterologous membrane protein production in Escherichia coli by combining transcriptional tuning and codon usage algorithms. PLoS One, 12, e0184355.

25. Clarke, T.F. and Clark, P.L. (2008) Rare codons cluster. PloS one, 3, e3412.

26. Wasels, F., Jean-Marie, J., Collas, F., López-Contreras, A.M. and Ferreira, N.L. (2017) A two-plasmid inducible CRISPR/Cas9 genome editing tool for Clostridium acetobutylicum. Journal of microbiological methods, 140, 5–11.

27. McAllister, K.N., Bouillaut, L., Kahn, J.N., Self, W.T. and Sorg, J.A. (2017) Using CRISPR-Cas9-mediated genome editing to generate C. difficile mutants defective in selenoproteins synthesis. Scientific reports, 7, 1–12.

28. Wang, S., Hong, W., Dong, S., Zhang, Z.-T., Zhang, J., Wang, L. and Wang, Y. (2018) Genome engineering of Clostridium difficile using the CRISPR-Cas9 system. Clinical Microbiology and Infection, 24, 1095–1099.

29. Wang, S., Dong, S., Wang, P., Tao, Y. and Wang, Y. (2017) Genome editing in Clostridium saccharoperbutylacetonicum N1-4 with the CRISPR-Cas9 system. Applied and environmental microbiology, 83, e00233–00217.

30. Nariya, H., Miyata, S., Kuwahara, T. and Okabe, A. (2011) Development and characterization of a xylose-inducible gene expression system for Clostridium perfringens. Applied and environmental microbiology, 77, 8439–8441.

31. Müh, U., Pannullo, A.G., Weiss, D.S. and Ellermeier, C.D. (2019) A xylose-inducible expression system and a CRISPR interference plasmid for targeted knockdown of gene expression in Clostridioides difficile. Journal of bacteriology, 201, e00711–00718.

32. Seo, S.O., Wang, Y., Lu, T., Jin, Y.S. and Blaschek, H.P. (2017) Characterization of a Clostridium beijerinckii spo0A mutant and its application for butyl butyrate production. Biotechnology and bioengineering, 114, 106–112.

33. Ravagnani, A., Jennert, K.C., Steiner, E., Grünberg, R., Jefferies, J.R., Wilkinson, S.R., Young, D.I., Tidswell, E.C., Brown, D.P. and Youngman, P. (2000) Spo0A directly controls the switch from acid to solvent production in solvent-forming clostridia. Molecular microbiology, 37, 1172–1185.

34. Creutzburg, S.C., Wu, W.Y., Mohanraju, P., Swartjes, T., Alkan, F., Gorodkin, J., Staals, R.H. and van der Oost, J. (2020) Good guide, bad guide: spacer sequence-dependent cleavage efficiency of Cas12a. Nucleic acids research, 48, 3228–3243.

35. Kim, H.K., Song, M., Lee, J., Menon, A.V., Jung, S., Kang, Y.-M., Choi, J.W., Woo, E., Koh, H.C. and Nam, J.-W. (2017) In vivo high-throughput profiling of CRISPR– Cpf1 activity. Nature methods, 14, 153–159.

36. Zhu, H. and Liang, C. (2019) CRISPR-DT: designing gRNAs for the CRISPR-Cpf1 system with improved target efficiency and specificity. Bioinformatics, 35, 2783–2789.

37. Liao, C., Ttofali, F., Slotkowski, R.A., Denny, S.R., Cecil, T.D., Leenay, R.T., Keung, A.J. and Beisel, C.L. (2019) Modular one-pot assembly of CRISPR arrays enables library generation and reveals factors influencing crRNA biogenesis. Nature communications, 10, 1–14.

38. Zadeh, J.N., Steenberg, C.D., Bois, J.S., Wolfe, B.R., Pierce, M.B., Khan, A.R., Dirks, R.M. and Pierce, N.A. (2011) NUPACK: analysis and design of nucleic acid systems. Journal of computational chemistry, 32, 170–173.

